# Sulfonated rhodamines as impermeable labelling substrates for cell surface protein visualization

**DOI:** 10.1101/2021.03.16.435698

**Authors:** Ramona Birke, Julia Ast, Dorien A. Roosen, Bettina Mathes, Kilian Roßmann, Christiane Huhn, Ben Jones, Martin Lehmann, Volker Haucke, David J. Hodson, Johannes Broichhagen

## Abstract

Sulfonated rhodamines that endow xanthene dyes with cellular impermeability are presented. We fuse charged sulfonates to red and far-red dyes to obtain Sulfo549 and Sulfo646, respectively, and further link these to SNAP- and Halo-tag substrates for protein self-labelling. Cellular impermeability is validated in live cell imaging experiments in transfected HEK cells and neurons derived from induced pluripotent stem cells (iPSCs). Lastly, we show that Sulfo646 is amenable to STED nanoscopy by recording membranes of SNAP/Halo-surface-labelled human iPSC-derived neuronal axons. We therefore provide an avenue for rendering dyes impermeable for exclusive extracellular visualization via self-labelling protein tags.

## INTRODUCTION

Fluorescent microscopy is often the method of choice to visualize and interrogate cell biology.^[1,2]^ Two major methods can be distinguished, that is the use of genetically-encoded fluorescent proteins or the use of small molecule fluorophores.^[2]^ The latter can be targeted by chemical fusion to a selective and tight small molecule binder, or by means of self-labelling protein tags.^[3–5]^ A plethora of fluorescent small molecules are available for microscopy, spanning different photophysical and chemical properties.^[6,7]^ Desirable properties are brightness, resistance to photobleaching, and cellular permeability.^[8–12]^ Depending on imaging modality, other properties might be desirable such as blinking or fluorogenicity. However, very few fluorescent dyes exist for exclusive SNAP- and Halo-tag labelling on cell surface proteins, best typified by transmembrane receptors. Within that repertoire, even fewer are suitable for stimulated emission depletion (STED) nanoscopy,^[13]^ since higher laser powers are required that may lead to photobleaching, although impermeable dyes were used in the beginning^[14]^. While robust and permeable dyes can be targeted to SNAP- or Halo-tag, non-specific background labelling may become problematic, especially in live cell applications.^[15]^ Moreover, this might be the case for cell surface receptor localization (Figure 1A), and as such it remains difficult to isolate surface and intracellular pools of receptors not only for nanoscopy.^[16]^ To restrict a priori fluorogenic dyes to the cell surface, we set out to synthesize rhodamines endowed with a cell impermeable sulfonate moiety. To achieve this, the rhodamine scaffold of fluorogenic and bright JaneliaFluor (JF) dyes^[8]^ was extended with a short sulfonate-containing linker to provide red and far-red colors (Sulfo549 and Sulfo646), which can be targeted to cell surface tags and receptors (Figure 1B). The versatility of this approach is highlighted in live cell imaging with various tagged constructs in different cell types including human induced pluripotent stem cells (iPSCs)-derived neurons, and in fixed neurons by STED nanoscopy to obtain high-definition resolution of axonal membranes.

**Figure 1:**
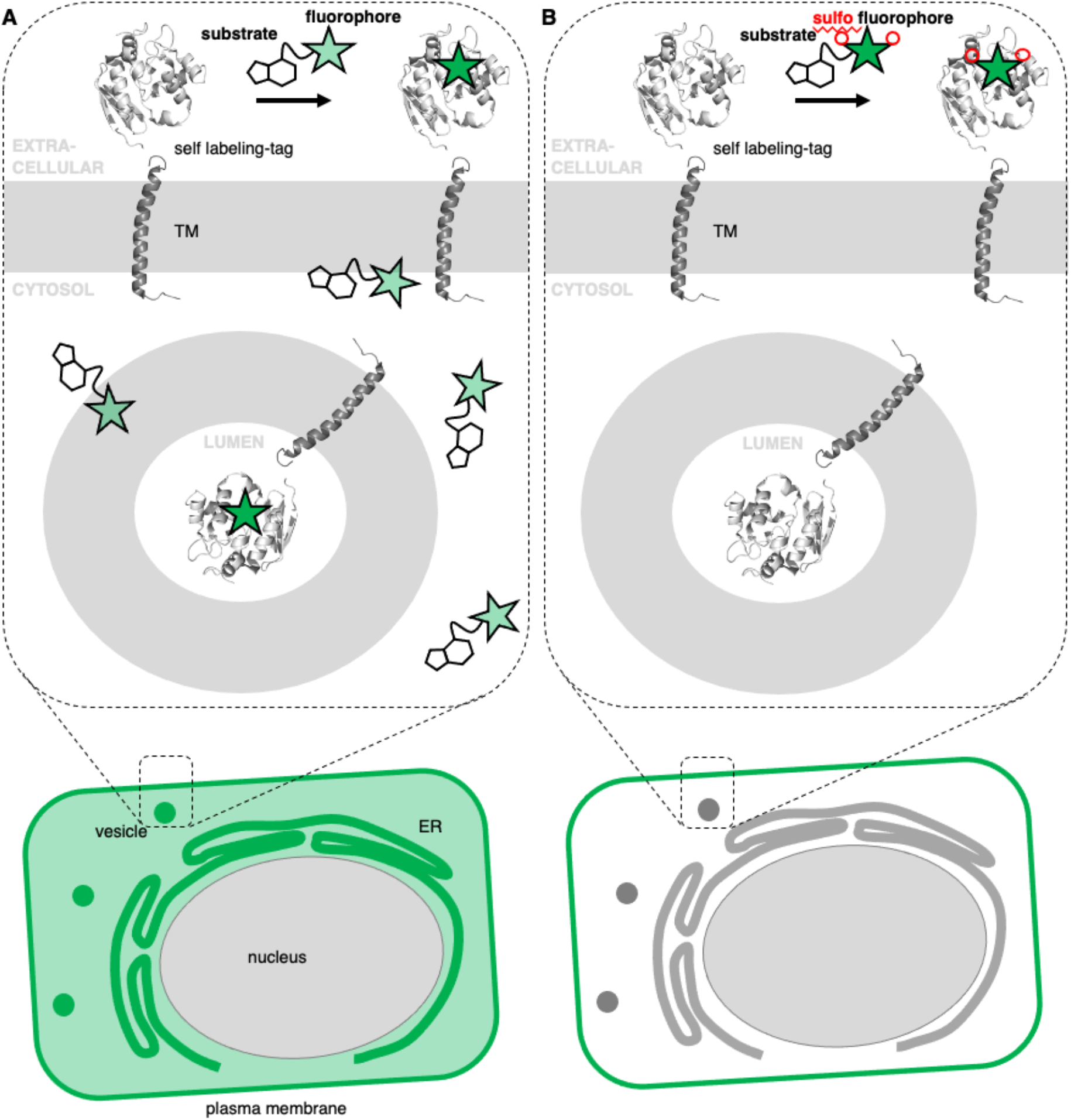
Permeable and impermeable fluorophores for protein labelling. A) A self-labelling tagged transmembrane protein is localized at the cell surface in addition to different intracellular, membrane-enclosed compartments (e.g. vesicles, ER) and is visualized by staining with a fluorescent dye. Unbound dyes lead to non-specific background signals. **B)** Sulfonation prevents the dye from entering the cell and therefore cleanly labels surface proteins, isolating them for visualization.

## RESULTS

We set out to synthesize impermeable dyes by the synthetic addition of charged sulfonates, which remain deprotonated and therefore not only become cell impermeable, but may also increase their solubility in aqueous media. With our primary aims in mind, i.e. i) impermeability, ii) labelling of SNAP- and Halo-tags, and iii) usage in STED nanoscopy, we decided to use xanthene dyes as a blueprint for our design, which are known to exist in two states, an open (fluorescent) and a closed (non-fluorescent) form (Figure 2A) and are among the most stable towards photobleaching. The recently reported JaneliaFluor dyes are rhodamine-based fluorophores, showing higher brightness and fluorogenicity than their tetramethyl rhodamines congeners, due to installment of azetidines as nitrogen containing moieties.^[8]^ As such, we synthesized Sulfo549 and Sulfo646 congeners by introducing a carboxylate handle on the 3-position of the azetidine, which was further derivatized to a sulfonated head group via peptide coupling to taurine (Figure 2A, Scheme S1). A carboxylate in the 6-position served as a position to install *O*^6^-benzylguanine (BG) or a chloroalkane (CA) group, which act as substrates for the self-labelling SNAP- and Halo-tag, respectively, and thereby obtained four molecules displaying two colors and two labelling modalities (Figure 2B, Scheme S1). In a first set of experiments, we assessed the excitation and emission profiles of our dyes in their unbound (i.e. BG- and CA-linked) and in their bound (i.e. SNAP- and Halo-tag reacted) states (Figure 2C). Fluorogenicity was reduced as expected when charges are added in close proximity to the dye, however, all dyes still showed high brightness (Figure S1, S2). Labelling was confirmed in vitro by incubation of Sulfo dyes with recombinant SNAP- and Halo-tag and subsequent mass spectrometry (see Supporting Information). Furthermore, we assessed kinetics of BG-Sulfo549 (*t*_1/2_ = 28.0 sec) labelling on SNAP-tag versus BG-TMR (*t*_1/2_ = 8.9 sec) and BG-JF_549_ (*t*_1/2_ = 15.3 sec) by means of fluorescent polarization, and found a slight decrease in rate of labelling by a factor of ~3.14 and ~1.83, respectively (Figure S3). Still, full labelling of SNAP:Sulfo549 was achieved within minutes and, interestingly, with enhanced polarization.

**Figure 2:**
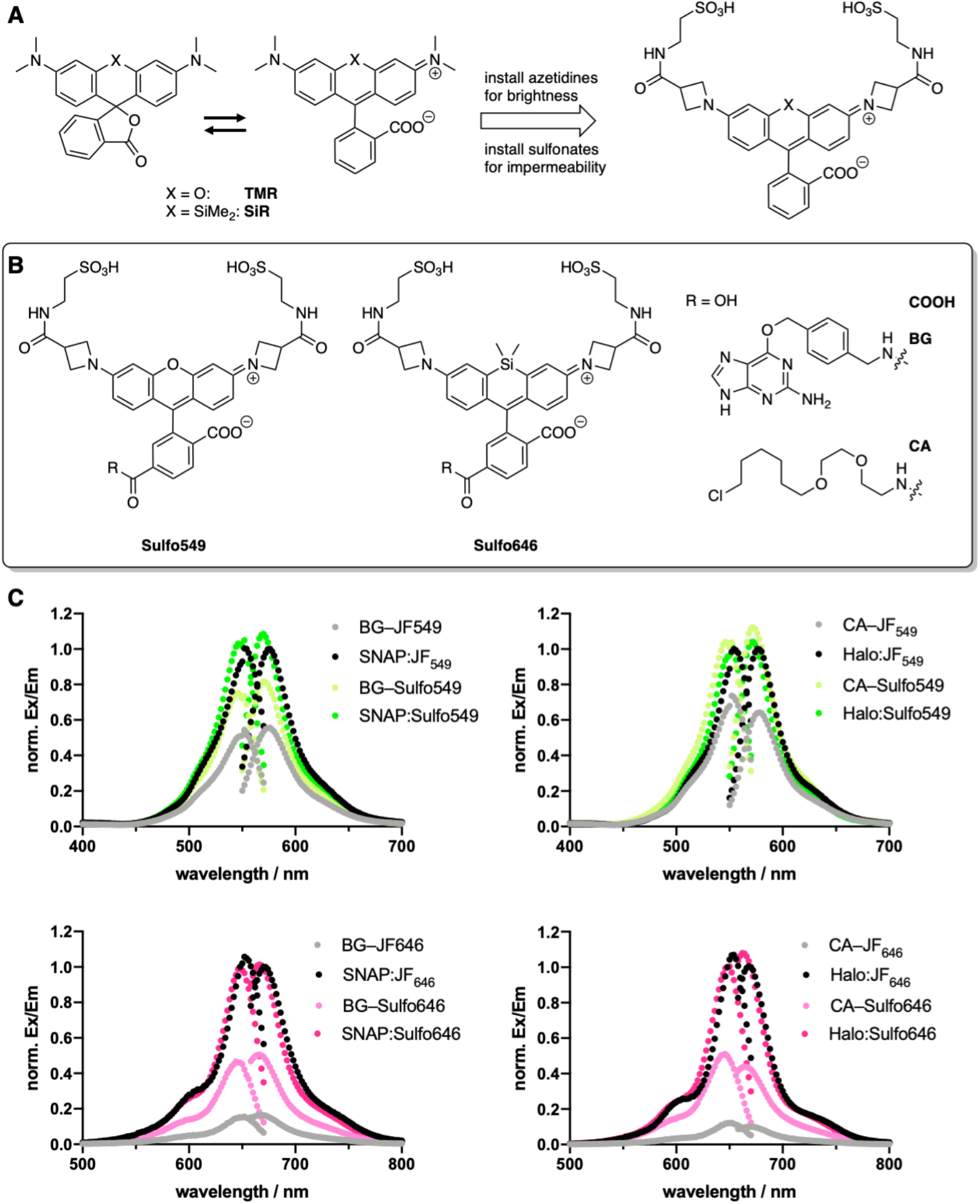
Design and properties of sulfonated rhodamines. **A)** Rhodamines display fluorogenic behavior between a closed, non-fluorescent form, and an open, fluorescent isomer. Azetidine installation enhances brightness, and can be further derivatized with sulfonates. **B)** Sulfo549 and Sulfo646 as red and far-red fluorophores that can be linked to BG and CA substrates for SNAP- and Halo-tag labelling, respectively. **C)** Normalized fluorescence excitation and emission spectra of non-sulfonated JF dyes and Sulfo dyes in solution and reacted with respective self-labelling tags.

Next, we wanted to test our molecules in a cellular setting for protein labelling in fluorescent microscopy. To prove that our impermeabilization strategy is applicable, we expressed a SNAP-Halo-construct with a nuclear localization signal (NLS; SNAP-Halo-NLS^[9]^) in HEK293T cells, before titrating 100-5000 nM of permeable JF_646_ or its impermeable counterpart Sulfo646. Clear concentration-dependent nuclear signals were detected for JaneliaFluor dyes, but not for the Sulfo probes (Figure S4). To confirm surface labelling, we cloned two constructs containing: i) an IgK trafficking signal for the plasma membrane; and ii) a SNAP-tag and Halo-tag separated by a single pass transmembrane (TM) domain (see Supporting Information). As such, the construct should be labelled exclusively on the surface when using impermeable dyes, while a permeable dye would lead to background staining from proteins residing in the cell. Indeed, by titration of 100-5000 nM of JF_646_ or Sulfo646 as before, we observed an increased background when using permeable dyes, and solely surface labelling when using Sulfo646, irrespective of the tag used (Figure S5).

Furthermore, due to the installation of two orthogonal tags on the same construct, we were able to dual-color label with a red and far-red dye, testing for permeability and localization properties. Having established that 100 nM of substrates lead to sufficient labelling, SNAP-TM-Halo-transfected HEK293T cells were incubated with a combination of BG-Sulfo646/CA-JF_549_ at each 100 nM concentration (Figure 3A). Widefield fluorescent imaging revealed surface localized staining for SNAP:Sulfo646 in combination with intracellular signals presumably stemming from non-surface trafficked or nascent Halo:JF_549_ (Figure 3B). This could be further resolved by plotting a line through a transfected cell (Figure 3C), which depicts plasma membrane and intracellular signals from the two colors. Using the same construct and settings, we then switched the dye colors for the respective tags. SNAP-TM-Halo transfected HEK293T cells were stained with BG-Sulfo549/CA-JF_646_ (Figure 3D), providing surface and intracellular staining for SNAP:Sulfo549 and Halo:JF_646_, respectively (Figure 3E), again shown using a line profile (Figure 3F). Finally, experiments were repeated by switching the labelling tag localization and transfecting HEK293 cells with Halo-TM-SNAP construct, this time labelled with BG-JF_646_/CA-Sulfo549 (Figure 4A). Again, widefield microscopy showed expected membrane associated signals for Halo:Sulfo549 and intracellular signals for SNAP:JF_646_ (Figure 4B, C). Using instead BG-JF_549_/CA-Sulfo646 (Figure 4D), we were able to switch the localization to external Halo:Sulfo646 and internal SNAP:JF_549_ (Figure 4E, F).

**Figure 3:**
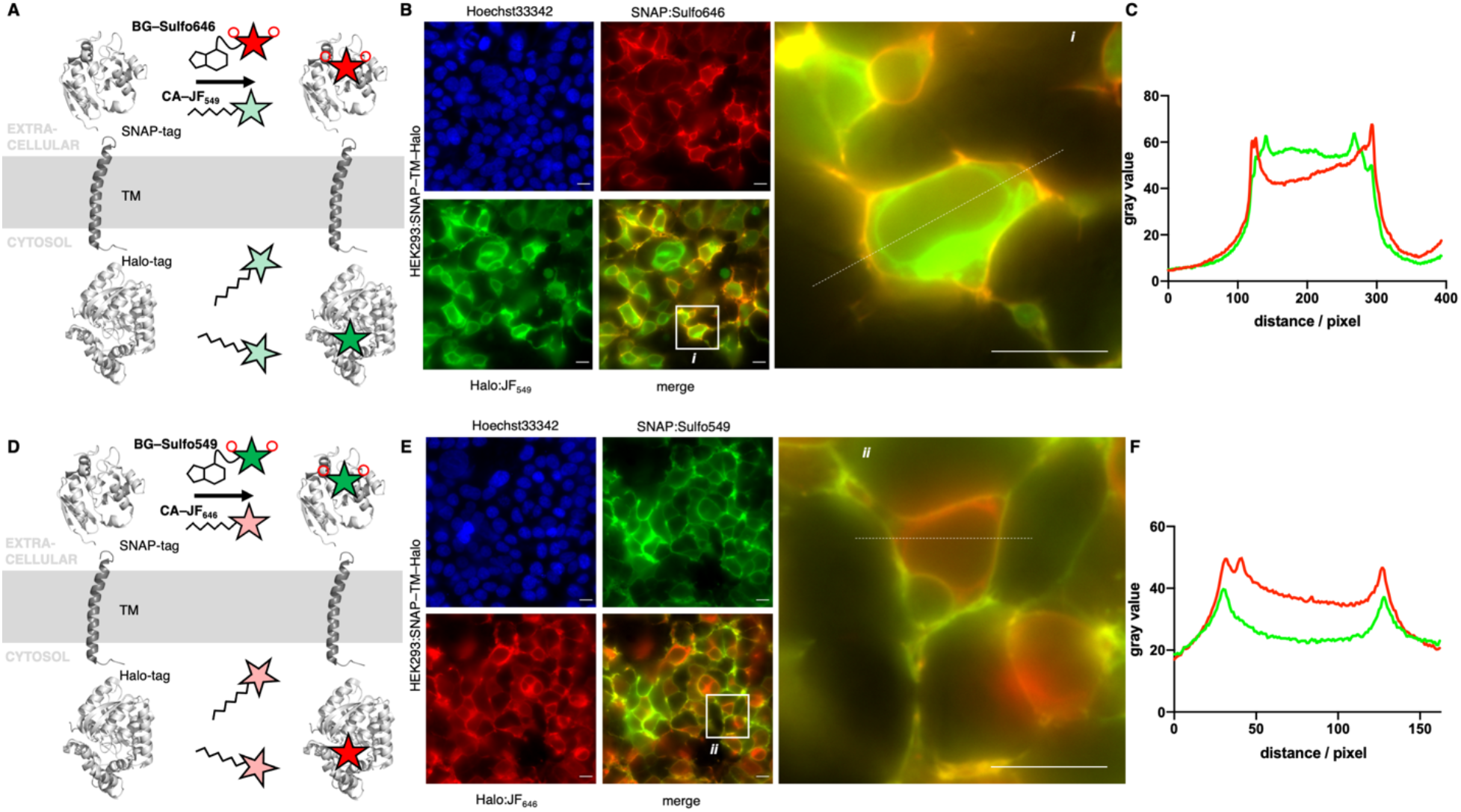
Widefield fluorescent imaging of live HEK293 cells transfected with SNAP-TM-Halo construct. **A)** Logic of labelling with BG-Sulfo646 and CA-JF_549_ leading to extra- and intracellular staining. **B)** Widefield imaging of live, transfected, and labelled HEK293T cells; insert shows zoom of one cell. **C)** Line plot intensity profile reveals membrane staining for impermeable Sulfo646 and intracellular labelling for JF_549_. **D-F)** As for A, B, C, but with BG-Sulfo549 and CA-JF_646_. Scale bar = 20 μm.

**Figure 4:**
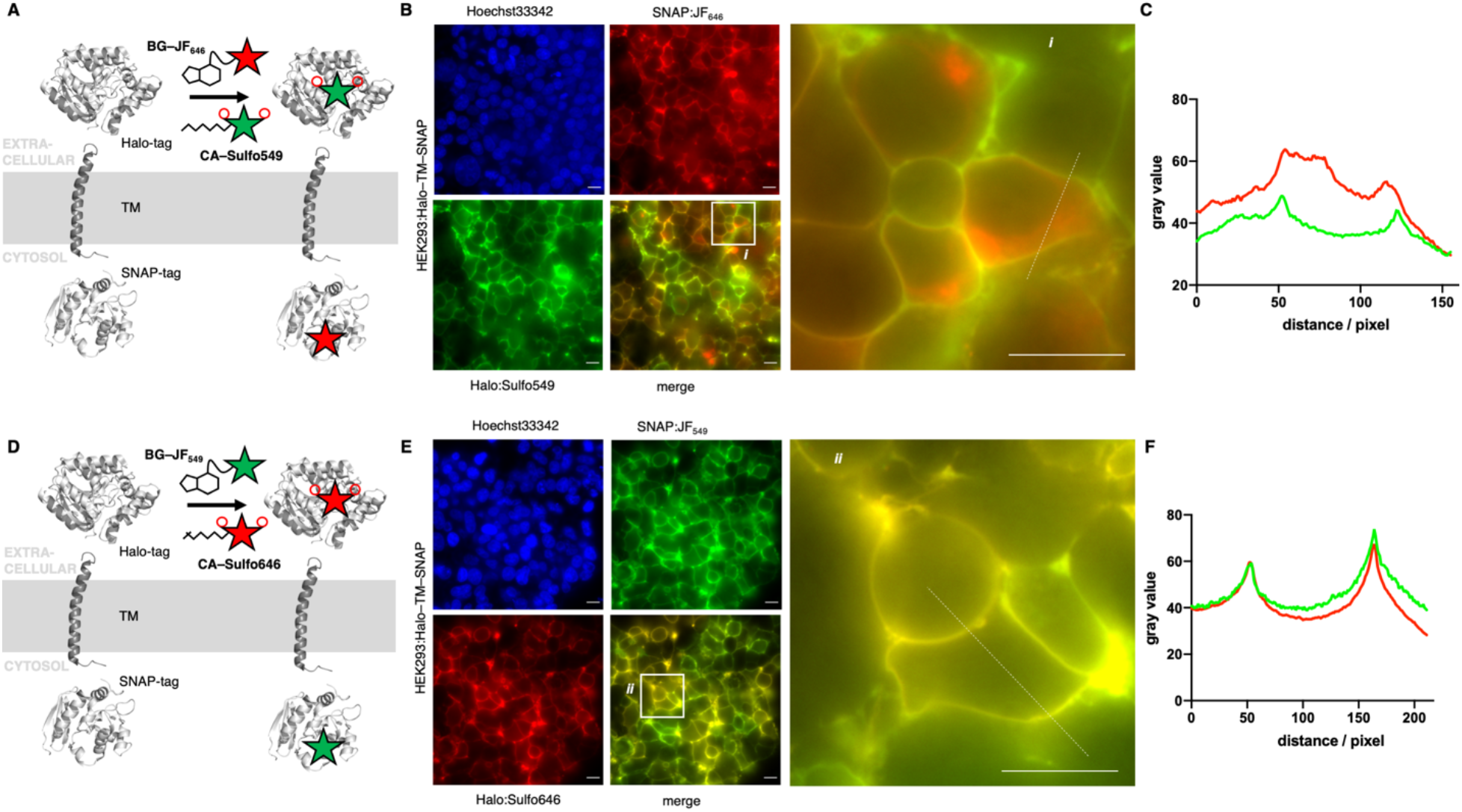
Widefield fluorescent imaging of live HEK293 cells transfected with Halo-TM-SNAP construct. **A)** Logic of labelling with CA-Sulfo549 and BG-JF_646_ leading to extra- and intracellular staining. **B)** Widefield imaging of live, transfected, and labelled HEK293T cells, insert shows zoom of one cell. **C)** Line plot intensity profile reveals membrane staining for impermeable Sulfo549 and intracellular labelling for JF_646_. **D-F)** As for A, B, C, but with CA-Sulfo646 and BG-JF_549_. Scale bar = 20 μm.

To show the utility of the Sulfo dyes for labelling cell surface receptors, we transfected AD293 cells with N-terminal SNAP- and Halo-tagged glucagon-like peptide-1 receptor (GLP1R), a class B GPCR and target for the incretin-mimetic class of anti-diabetic therapy (Figure 5). In its non-stimulated state, GLP1R displays minimal constitutive activity and is largely present at the cell surface. Notably, differences between JF_549_ and JF_646_ and their Sulfo derivatives were present for SNAP labelling, and became even more prominent for Halo labeling, where a large improvement in membrane resolution and brightness was detected.

**Figure 5:**
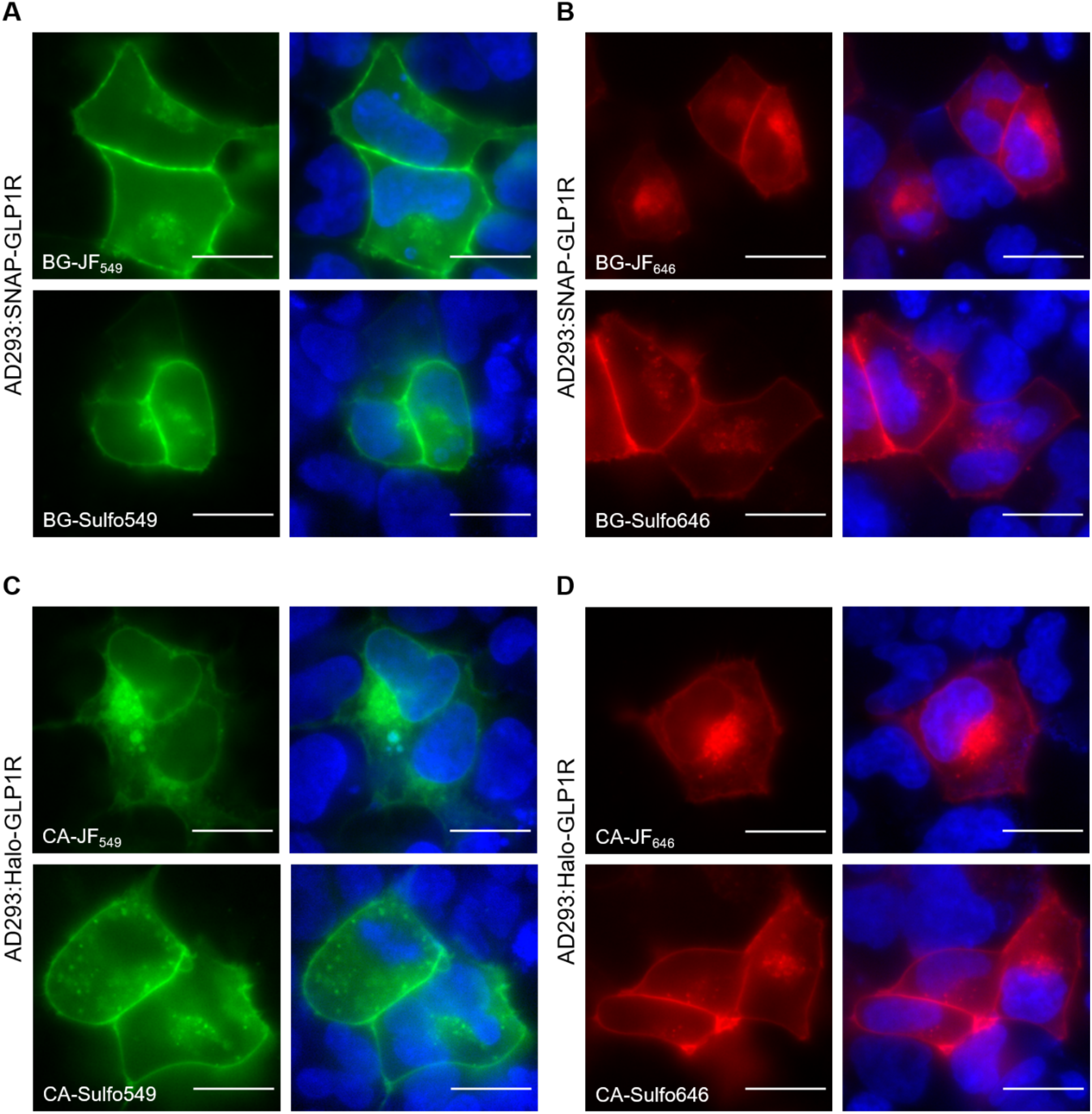
SNAP/Halo-GLP1R labelling. **A-B)** Widefield imaging of AD293 cells expressing SNAP-GLP1R labeled with BG-JF_549_, BG-Sulfo549, BG-JF_646_, or BG-Sulfo646, and Hoechst33342 staining. **C-D)** Same as for A and B but AD293 expressing Halo-GLP1R and labeled with CA versions of dyes. Scale bar = 20 μm.

Given the performance of the Sulfo dyes so far in heterologous cell systems, we sought to extend studies to more complex cell populations where background signal can make accurate protein localization difficult. To allow this, human cortical neurons were derived from iPSC and co-cultured with murine primary astrocytes^[17,18]^, before transfection with our SNAP-TM-Halo and Halo-TM-SNAP constructs. Labeling was performed with the respective impermeable far-red Sulfo646 and permeable red JF_549_, before live imaging was conducted by confocal microscopy.

Using the SNAP-TM-Halo construct labelled with BG-Sulfo646 and CA-JF_549_, clear labelling was observed for BG-Sulfo646 (Figure 6A). Likewise, Halo-TM-SNAP labelled with CA-Sulfo646 and BG-JF_549_ (Figure 6B) showed clearest labelling for CA-Sulfo646. Thus, for both self-labelling tags, the Sulfo dyes demonstrated excellent performance for cell surface protein visualization in more complex cell types.

**Figure 6:**
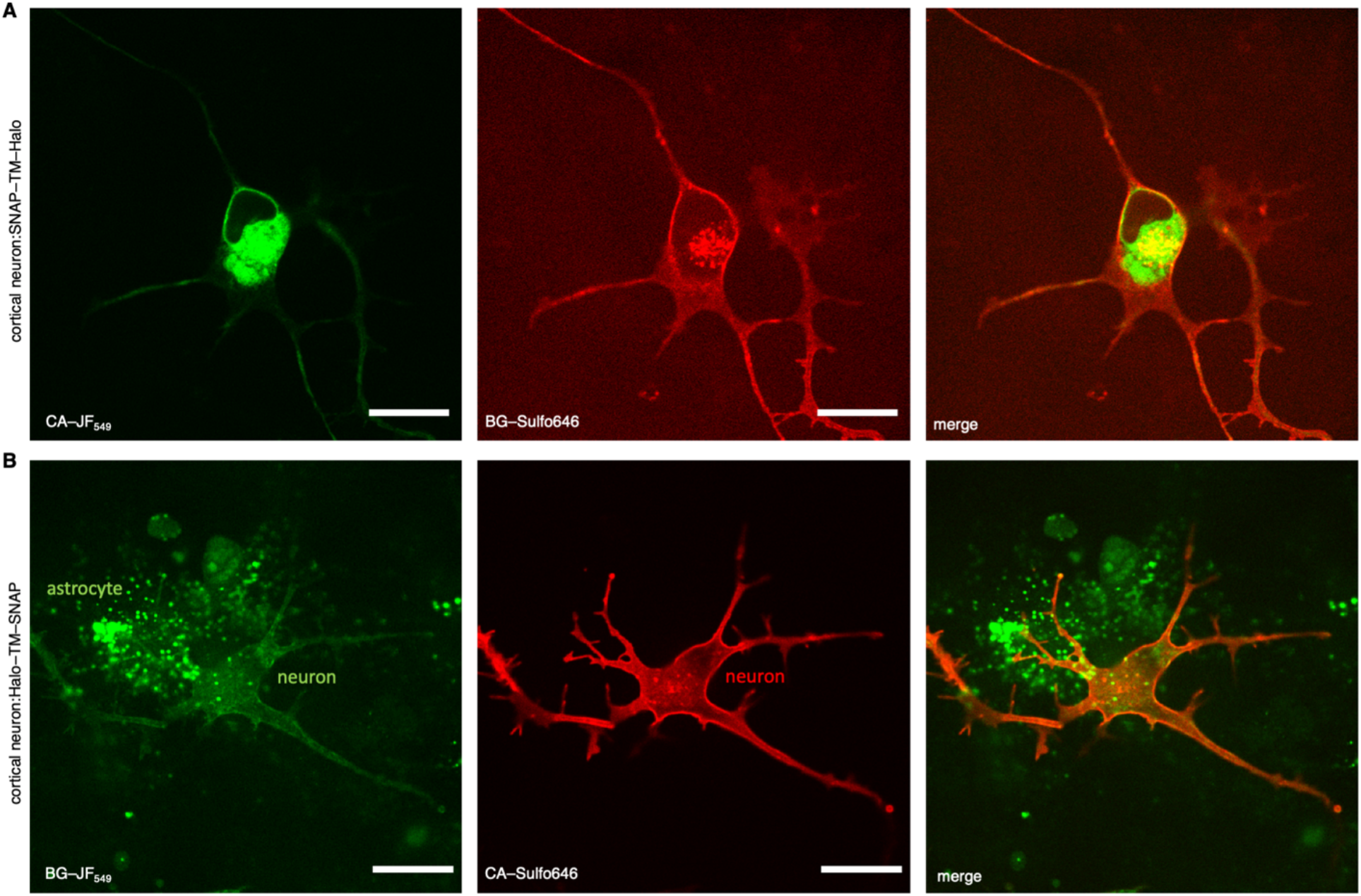
Confocal fluorescence imaging of live transfected human cortical neurons. Neurons co-cultured with astrocytes were transfected with a SNAP-TM-Halo (A) or a Halo-TM-SNAP (B) construct and labelled with CA-JF_549_/BG-Sulfo646 (A) and BG-JF_549_/CA-Sulfo646. Scale bar = 20 μm.

Lastly, we fixed the cultures before STED nanoscopy on Halo:Sulfo646 and SNAP:JF_549_ labelled neurons (Figure 7A). An accumulated line profile along an axon (white box) of the confocal and STED images revealed that Sulfo646 is amenable to nanoscopy (Figure 7B). By two-Gaussian fitting, we obtained sharper full width half-maximal values for STED versus confocal (FWHM_confocal_ = 268.5 and 292.3 nm; FWHM_STED_ = 157.5 and 223.1 nm) microscopy. Consistent with our previous results, Halo:Sulfo646 generated a more pronounced signal along the axonal membrane, while SNAP:JF_549_ could be observed in the intra-axonal compartment (Figure 7C).

**Figure 7:**
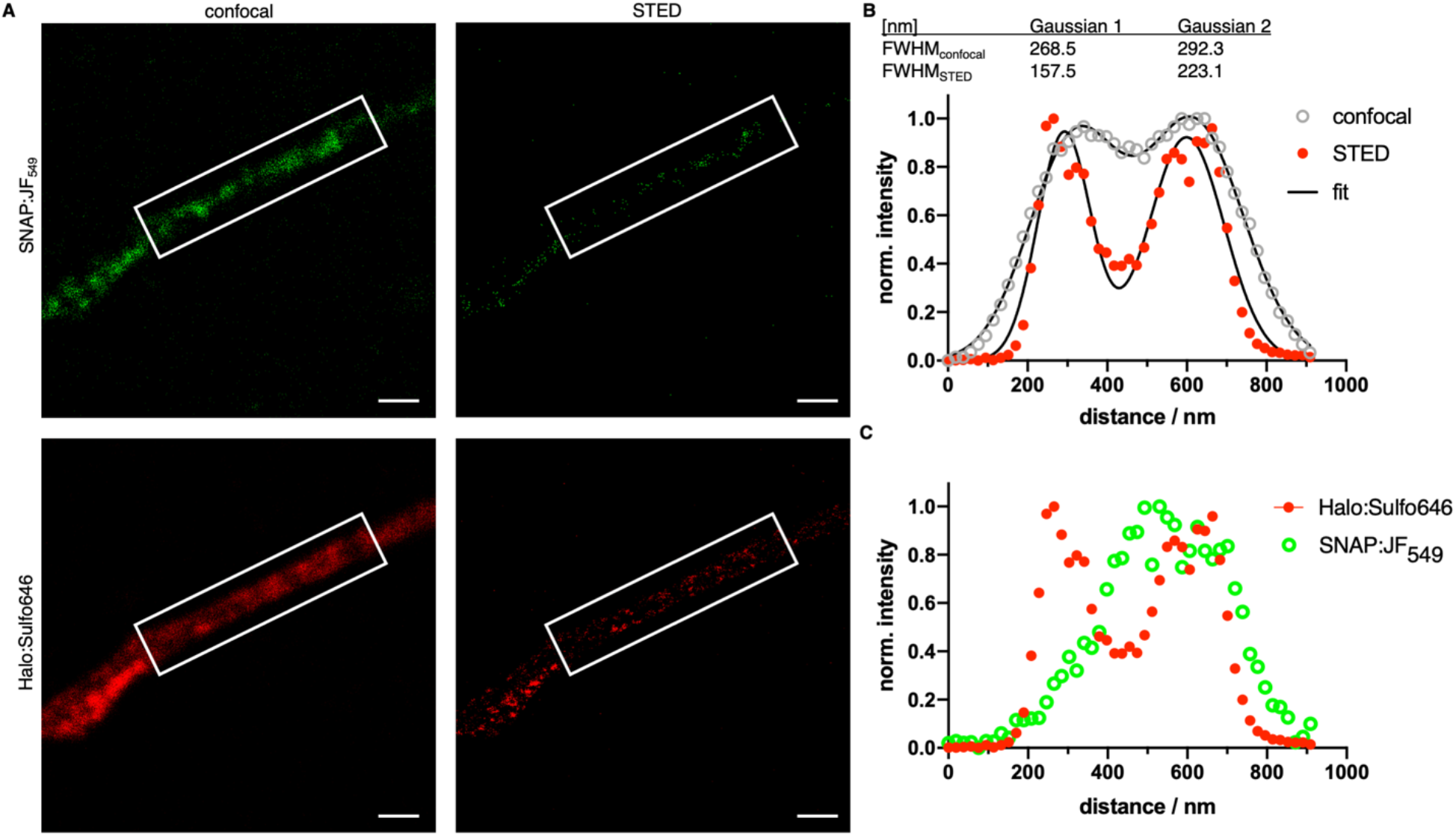
Confocal and STED imaging of fixed human cortical neurons. **A)** Cultures transfected with Halo-TM-SNAP from Figure 6 were fixed and BG-JF_549_- and CA-Sulfo646-labelled axons imaged by confocal microscopy and STED nanoscopy. **B)** Line profile along an axon showing improved resolution of Halo:Sulfo646 at the membranous margins in STED versus confocal imaging. **C)** Halo:Sulfo646 allows visualization of surface signals, whereas SNAP:JF_549_ is largely confined to the intra-axonal compartment (STED signals). Scale bar = 1 μm.

## DISCUSSION

The need for custom-tailored dyes is in high demand as the range of microscopy modalities, experimental techniques and labelling strategies increases. Recent developments^[9,19–22]^ have focused mostly on boosting brightness, fluorescent lifetimes, chemical stability and/or fluorogenicity, the latter being a cause for cellular permeability. For the interrogation of cell surface proteins that are genetically fused to self-labelling protein tags (e.g. SNAP- and Halo-tag), however, cell impermeable dyes are desirable. Rendering dyes impermeable is usually achieved by introduction of sulfonates, which remain negatively charged in biological systems and are therefore not able to cross the plasma lipid bilayer. While many sulfonated dyes exist, such as Alexa488/568/647, or LD555/655^[23]^, their application for enzyme self-labelling and STED nanoscopy has so far not reached the performance bar set by permeable dyes. A recent study, however, showed great performance of LD dyes for FRET measurements of dimer formation on SNAP-tagged GPCRs and in TIRF microscopy for single molecule FRET recovery after photobleaching.^[24]^ Rhodamine dyes have attracted some attention in recent years since the emergence of JaneliaFluor dyes, which rely on the exchange of azetidine groups for dimethylamines on various molecular scaffolds^[8]^. Indeed, one impermeable version has been described, JF_635i_, which retains some of its fluorogenicity and has been used to observe Halo-tagged transferrin receptor recycling.^[25]^ Nevertheless, it has only been described as a Halo-tag substrate and has not been subjected to super-resolution STED nanoscopy. We aimed to expand this palette, by using azetidine containing rhodamines for red and far-red imaging, which show minimal fluorogenicity and therefore maximal brightness. Accordingly, we synthesized Sulfo549 and Sulfo646 based on JF_549_ and JF_646_, each bearing two sulfonate groups, and further linked them to SNAP- and Halo-tag substrates BG and CA, respectively. Our probes add favorably to a previous study, where we reported on a strategy to limit any dye to cell surface exposed SNAP tags by altering the BG substrate to a sulfonate itself (termed SBG)^[16]^. With a simple chloride anion being the leaving group for the Halo-tag, introducing a charged sulfonate is not tolerated. Therefore, the impermeable characteristics need to instead be provided by the dye, for which we provide a solution herein.

To test our approach, we cloned constructs for cellular transfections that bear a SNAP- and Halo-tag separated by a transmembrane domain, thereby placing each tag either side of the plasma membrane. These constructs allowed screening of permeability parameters in live HEK293 cells. Sulfo dyes performed well in these systems, where no background signals from intracellular spaces were detected. Line scans revealed clean plasma membrane staining of Sulfo dyes and intracellular staining of the permeable JF dyes.

Encouraged by this, we next used SNAP- and Halo-tagged GLP1R to provide performance benchmarking in a more relevant cell surface signaling molecule. Comparable to previous findings with the SNAP/Halo constructs, we obtained clean surface labelling for both colors and both protein tags, with more prominent effects for Halo, which is in line with in vitro measurements. Given that GLP1R labelling was performed at 37 °C, where constitutive activity of GPCRs can be increased, intracellular signal from Sulfo dyes can be observed, and we note that their strength might differ due to different expression and endocytosis levels. While we have shown dual color GLP1R trafficking in a previous report on a SNAP-tagged construct^[16]^, our results herein set the stage for flexible use of colors on the Halo-tag not only on GLP1R, for instance for post endocytic protein trafficking^[25,26]^ and the interrogation of cell surface receptor ensembles^[27–29]^.

We were also able to extend this approach to more complex cell types, by staining live somatodendritic and axonal compartments in human cortical neurons derived from iPSC, co-cultured with astrocytes. By transfecting neurons with SNAP-TM-Halo or Halo-TM-SNAP, we aimed to benchmark far-red staining of the outer membrane with both tags, as this color was our choice for later super-resolution imaging. As such, we observed surface labelling with both constructs and the use of the respective Sulfo646, with signals markedly brighter when bound to Halo. In contrast, CA-JF_549_ clearly stained the intracellular Halo-tagged protein pool of neurons, while BG-JF_549_ non-specifically accumulated in co-cultured astrocytes. Self-labelling tags have been employed in “brainbow” labelling, where they offer more flexibility in terms of colors available, more straightforward applicability than antibodies, and better survival of the rather harsh clearing conditions with respect to fluorescent proteins.^[30]^ Indeed, we anticipate Sulfo dyes to be a favorable addition to such studies and their performance in whole tissues^[31,32]^, and with this observed trend in mind, we chose to continue with Halo:Sulfo646 in our preparations.

For this reason, we fixed the astrocyte/neuron co-culture and tested SNAP:JF_549_ and Halo:Sulfo646 for STED nanoscopy. As expected, JF_549_ was not amenable to the depletion laser, but gratifyingly, Sulfo646 was able to improve full width half maximal values when a broad line plot was applied along an axon in STED imaging. In addition, the resolution, i.e. the distance of the two maxima, of the membranes were resolved to be ~300 nm in both confocal and STED. Previous studies resolved the median axonal diameter of organotypic GFP expressing CA1 neurons to be 203 nm by STED^[33]^, and to be 242 nm by super-resolution shadow imaging (SUSHI)^[34]^.

## SUMMARY

We have designed and synthesized sulfonated fluorescent rhodamines dyes (Sulfo549 and Sulfo646) that are based on the JaneliaFluor scaffolds to obtain bright and impermeable dyes in the red and far-red. By linking these dyes to substrates recognized by the SNAP and Halo-tag, we were able to achieve exclusive cell surface labelling in HEK293/AD293 cells and in human induced neurons by means of widefield and confocal microscopy. Lastly, we employed STED nanoscopy on Sulfo646-labelled neuron axons and showcase their performance by resolving the axonal membranes. We anticipate that these and other sulfonated rhodamines will be useful for visualizing cell surface proteins using a range of imaging approaches spanning widefield through super-resolution.

## METHODS

### Chemistry, cloning and in vitro protein labelling

Chemical Schemes, synthetic protocols, protein labelling in vitro, and characterization can be found in the Supporting Information. Plasmids were cloned using Gibson assembly cloning Kit (NEB), primer were designed using the NEBuilder assembly tool. Plasmids were isolated using a mini prep kit (Thermo Fisher). DNA concentration was measured on a NanoDrop (Thermo Fisher) and verified by Sanger sequencing (see Supporting Information).

### In vitro fluorescence spectroscopy

Purified SNAP_f_ and Halo was obtained as previously described.^[20]^ Labelling dyes were dissolved in DMSO to a concentration of 1 mM and diluted in activity buffer (containing: 50 mM NaCl, 50 mM HEPES, pH=7.3 + 4 μg/mL BSA) to 500 nM. Protein was diluted in activity buffer to a concentration of 2 μM. 100 μL of each protein and labelling agent were combined in each well in a black flat bottom 96-well plate and allowed to incubate at r.t. for 30 min, before fluorescence spectra were acquired in quadruplicates on a TECAN infinite 2000Pro plate reader. Experiments were run in quadruplicate and plotted in GraphPad Prism 8.

Kinetic measurements were performed as previously described on a TECAN GENios Pro plate reader by means of fluorescence polarization. Stocks of SNAP_f_ (2 μM) and substrates (200 nM) were prepared in activity buffer (containing in mM: NaCl 50, HEPES 50, pH 7.3) with additional 10 μg/mL BSA. SNAP_f_ and substrates were mixed (100 μL each) in a Greiner black flat bottom 96 well plate. Mixing was performed *via* a built-in injector on a TECAN GENios Pro. Fluorescence polarization reading was started immediately (λ_Ex_ = 535±25 nm; λ_Em_ = 590±35 nm; 10 flashes; 40 μs integration time). Experiments were run in quadruplicates and raw polarization values were one-phase decay fitted in GraphPad Prism 8.

### Protein mass spectrometry

Labelling substrates were dissolved in DMSO to a concentration of 1 mM and diluted in activity buffer (containing: 50 mM NaCl, 50 mM HEPES, pH=7.3 + 4 μg/mL BSA) to 20 μM. Protein was diluted in activity buffer to a concentration of 2 μM. 25 μL of each protein and labelling agent were combined in a mass spec vial and allowed to incubate at r.t. for 1h, before full protein mass was acquired. In case for non-labelling control, 25 μL of activity buffer was mixed with 25 μL of protein.

### Cell culture, staining and microscopy

#### HEK293

HEK293T cells were cultured in growth medium (DMEM, Glutamax, 4.5 g Glucose, 10% FCS, 1% PS; Invitrogen) at 37 °C and 5% CO_2_. 30,000 cells/well were seeded on 8-well μl slides (Ibidi) previously coated with 0.25 mg/ml poly-L-lysine (Aldrich, mol wt 70000-150000). The next day, 400 ng DNA was transfected using 0.8 μl Jet Prime reagent in 40 μl Jet Prime buffer (VWR) per well. Medium was exchanged against antibiotic free media before the transfection mix was pipetted on the cells. After 4 hours incubation at 37 °C and 5% CO_2_, medium was exchanged against growth media. After 24 hours cells were stained and imaged. All dyes were used in a concentration of 100 nM. 5 μM Hoechst 33342 was used to stain DNA. Staining was done in growth medium at 37 °C, 5% CO_2_ for 30 minutes. Afterwards cells were washed once in growth media and imaged live in cell imaging buffer (Invitrogen) using an Epifluorescence Microscope, TIE Nikon equipped with pE4000 (cool LED), Penta Cube (AHF 66-615), 60x oil NA 1.49 (Apo TIRF Nikon) and imaged on sCMOS camera (Prime 95B, Photometrics) operated by NIS Elements (Nikon). For excitation the following LED wavelengths were used: Hoechst – 405 nm, JF_549_ and Sulfo549 – 550 nm, JF_646_ and Sulfo646 – 635 nm.

#### AD293

AD293 cells were cultured in DMEM (D6546, Merck) supplemented with 10% FCS (Merck) and 2 mM L-glutamine (Thermo Scientific) at 37 °C and 5% CO_2_. Cells were seeded on poly-L-lysine-coated (MW > 300,000, 0.1 mg/ml, Biochrom) 8-well chamber slides (Nunc Lab-Tek II) and transfected the next day (50-70% confluency). For transfection, growth media was exchanged against Opti-MEM (Gibco), and 110 ng plasmid DNA (SNAP-GLP1R from cisbio, for sequence of Halo-GLP1R see SI) and 0.3 μl Lipofectamine 2000 (Invitrogen) per chamber were mixed in Opti-MEM, incubated for five minutes and then added to the cells. After incubation at 37 °C and 5% CO_2_ for 5 hours, Opti-MEM was exchanged against growth media. AD293 cells were labeled and imaged 24 h after transfection. Cells were labelled in growth media containing 500 nM of dyes for 30 minutes at 37 °C and 5% CO_2_. For the last five minutes of incubation, Hoechst33342 (4.4 μM) was added. After one wash, cells were imaged in growth media using a Crest X-Light spinning disk head coupled to a Nikon Ti-E automated base and 60x/1.4 NA objective. Excitation was delivered at *λ* = 395/25 nm (Hoechst), 575/25 nm (JF_549_ and Sulfo549), and 640/30 nm (JF_646_ and Sulfo646) using a Lumencor Spectra X Light engine, and emitted signals were detected at *λ* = 460/50 nm, 630/75 nm, and 700/75 nm, respectively, using a Photometrics Evolve Delta 512 EMCCD.

#### Human iPSC-derived neurons

Human induced pluripotent stem cells (iPSCs) engineered to express mNGN2 under a doxycycline-inducible system in the AAVS1 safe harbor locus were used for the i^3^Neuron differentiation protocol, as described previously.^[17,18]^ In brief, iPSCs were seeded on Matrigel (Corning)-coated dishes in StemFlex Medium (Gibco) supplemented with 5 nM Y-27632 dihydrochloride ROCK inhibitor (Stem Cell Technologies). The iPSCs were subsequently fed daily for 3 consecutive days with Neuronal Induction Medium (DMEM/F12 medium (Gibco) containing 2.5 μg/mL doxycycline, 1X N2‐supplement (Gibco), 1X NEAA (Gibco), 1X GlutaMAX (Gibco), 1X Pen/Strep (Gibco), 10 ng/mL BDNF (PeproTech), 10 ng/mL NT‐3 (PeproTech) and 1 μg/mL Laminin (Gibco)). On the third day, Neuronal Induction Medium was supplemented with 6 μg/ml puromycin. After 3 days, pre-differentiated i^3^Neurons were dispersed using Trypsin (Gibco) and co-cultured with primary murine astrocytes on matrigel-coated 25 mm glass coverslips in Neuron Culture Medium (NeuroBasal Medium (Gibco), supplemented with 1X B27 (Gibco), 1X GlutaMAX (Gibco), 1X Pen/Strep (Gibco), 10 ng/mL BDNF (PeproTech), 10 ng/mL NT‐3 (PeproTech) and 1 μg/mL Laminin (Gibco)). 50% of the Neuron Culture Medium was replaced every 2-3 days and supplemented with 2 μM araC 5 days after culturing, to limit glial proliferation. i^3^Neurons were transfected, labeled and imaged as described below 10-12 days after co-culturing.

#### Neuronal transfection, labeling and imaging

i^3^Neurons were transfected with 2 μg of plasmid (SNAP-TM-Halo or Halo-TM-SNAP) per 25 mm coverslip using a calcium phosphate transfection kit (Promega), according to manufacturer’s instruction. 24 hours after transfection, cells were labeled with 500 nM in parallel of either BG-Sulfo646 and Halo-JF_549_, or Halo-Sulfo646 and BG-JF_549_, dissolved in Neuron Culture Medium for 30 minutes at 37 °C.

For live imaging, i^3^Neurons were washed once with Neuron Culture Medium before imaging in conditioned Neuron Culture Medium using a spinning disc confocal microscope (Ti Eclipse, Nikon) equipped with a spinning disk (CSU-X1, Yokogawa), EMCCD Camera (AU-888, Andor), 60x Plan-Apo NA 1.40 objective (oil immersion, Nikon), incubation chamber (37 °C, 5% CO_2_, Okolab). JF_549_ and Sulfo646 were excited with 561 nm and 638 nm laser, respectively, and emission was detected within 600-650 and 700-775 nm filter range, respectively. Images were taken using an additional 1x lens, resulting in 110 nm effective pixel size.

For super-resolution imaging, labelled i^3^Neurons were washed once with PBS before fixing 20 minutes at room temperature with 4% PFA and 4% sucrose in PBS. Fixation solution was removed and fixed i^3^Neurons were incubated with quenching solution (0.1 M glycine, 0.1 M NH_4_Cl in PBS) for 10 minutes at room temperature. Coverslips were subsequently washed once with PBS and once with water, mounted in ProLong Gold Antifade (ThermoFisher), and cured for 24 h at room temperature. STED imaging was performed on a Leica TCS 3x gSTED microscope, equipped with a pulsed white light excitation laser (NKT Photonics) and a 775 nm depletion laser. Two-channel STED imaging was performed by sequentially exciting JF_549_ or Sulfo646 at 550 and 640 nm, respectively, using a 100x PL Apo NA1.4 objective (oil immersion, Leica). The 775 nm STED laser was used to deplete both JF_549_ and Sulfo646. Time-gated detection was set from 0.5-6 ns for all dyes and emission was detected within 560-643 and 650-751 nm, respectively. Fluorescence signal was detected sequentially by two hybrid detectors, 6-fold zoom, 8-bit sampling and 1,024×1,024 pixel scanning format, resulting in 18.9×18.9 nm pixel dimension.

## ETHICAL STATEMENT

The human iPSC cell line (BIHi005-A) was generated following authorisation by the donor and ethics approval from the primary project and has been registered in the European Human Pluripotent Stem Cell Registry (hPSCreg): http://hpscreg.eu/cell-line/BIHi005-A. Cells were kindly provided by Dr. Sebastian Diecke, MDC, Berlin, Germany. Preparation of primary murine astrocytes (C57BL/6) was reviewed and approved by the ethics committee of the “Landesamt für Gesundheit und Soziales” (LAGeSo) Berlin and were conducted accordingly to the committee’s guidelines.

## AUTHOR CONTRIBUTIONS

JB designed and conceptualized the study. BM, KR, CH, and JB performed chemical synthesis and characterization. JA, DR, ML, and JB performed microscopy. BJ provided reagents. ML, VH, DJH, and JB supervised the study. DJH and JB wrote the manuscript with input from all the authors.

## ACKNOWLEDGEMENTS

Supported by grants from the Deutsche Forschungsgemeinschaft (DFG, German Research Foundation, GRK2318/TJ-Train A4 to ML and VH and HA2686/19-1 - NeuroNex2 to V.H.). BJ acknowledges support from the Academy of Medical Sciences, Society for Endocrinology, The British Society for Neuroendocrinology, the European Federation for the Study of Diabetes, and an EPSRC capital award. DJH was supported by MRC (MR/N00275X/1 and MR/S025618/1) and Diabetes UK (17/0005681) Project Grants. This project has received funding from the European Research Council (ERC) under the European Union’s Horizon 2020 research and innovation programme (Starting Grant 715884 to DJH and Advanced Grant 884281 to VH). We thank Sarah Mikami for synthetic assistance, Cornelia Ulrich and Sebastian Fabritz for mass spectrometry, Kai Johnsson for providing SNAP- and Halo-tag containing plasmids, chemical precursors and constant support (all MPIMR).

