## Supporting Information for "Sulfonated rhodamines as impermeable labelling substrates for cell surface protein visualization"

[1] Leibniz-Forschungsinstitut für Molekulare Pharmakologie, Berlin, Germany.

[2] Institute of Metabolism and Systems Research (IMSR), University of Birmingham, Birmingham, B15 2TT, UK.

[3] Centre for Endocrinology, Diabetes and Metabolism, Birmingham Health Partners, Birmingham, B15 2TT, UK.

[4] Centre of Membrane Proteins and Receptors (COMPARE), University of Birmingham, Birmingham, UK.

[5] Department Molecular Pharmacology and Cell Biology, Leibniz-Forschungsinstitut für Molekulare Pharmakologie, Berlin, Germany.

[6] Department of Chemical Biology, Max Planck Institute for Medical Research, Heidelberg, Germany.

[7] Section of Endocrinology and Investigative Medicine, Imperial College London, London W12 0NN, United Kingdom.

**Table of Contents**

**1. General** 3

**2. Synthesis** 4

*2.1. General procedure A for Buchwald-Hartwig coupling and deprotection* 4

*2.2. General procedure B for taurine coupling and deprotection* 4

*2.3. General procedure C for peptide coupling with TSTU* 4

*2.4. 4-(tert-Butoxycarbonyl)-2-(3-(3-carboxyazetidin-1-ium-1-ylidene)-6-(3-carboxyazetidin-1-yl)-3H-xanthen-9-yl)benzoate (2a)* 5

*2.5. 4-Carboxy-2-(3-(3-((2-sulfoethyl)carbamoyl)azetidin-1-ium-1-ylidene)-6-(3-((2-sulfoethyl)carbamoyl)azetidin-1-yl)-3H-xanthen-9-yl)benzoate (3a)* 5

*2.6. 4-((4-(((2-Amino-9H-purin-6-yl)oxy)methyl)benzyl)carbamoyl)-2-(3-(3-((2-sulfoethyl)carbamoyl)azetidin-1-ium-1-ylidene)-6-(3-((2-sulfoethyl)carbamoyl)azetidin-1-yl)-3H-xanthen-9-yl)benzoate (BG-Sulfo549)* 5

*2.7. 4-((2-(2-((6-Chlorohexyl)oxy)ethoxy)ethyl)carbamoyl)-2-(3-(3-((2-sulfoethyl)carbamoyl)azetidin-1-ium-1-ylidene)-6-(3-((2-sulfoethyl)carbamoyl)azetidin-1-yl)-3H-xanthen-9-yl)benzoate (Halo-Sulfo549)* 6

*2.8. 4-(tert-Butoxycarbonyl)-2-(3-(3-carboxyazetidin-1-ium-1-ylidene)-7-(3-carboxyazetidin-1-yl)-5,5-dimethyl-3,5-dihydrodibenzo[b,e]silin-10-yl)benzoate (2c)* 6

*2.9. 4-Carboxy-2-(5,5-dimethyl-3-(3-((2-sulfoethyl)carbamoyl)azetidin-1-ium-1-ylidene)-7-(3-((2-sulfoethyl)carbamoyl)azetidin-1-yl)-3,5-dihydrodibenzo[b,e]silin-10-yl)benzoate (3c)* 6

*2.10. 4-((4-(2-(2-Amino-9H-purin-6-yl)ethyl)benzyl)carbamoyl)-2-(5,5-dimethyl-3-(3-((2-sulfoethyl)carbamoyl)azetidin-1-ium-1-ylidene)-7-(3-((2-sulfoethyl)carbamoyl)azetidin-1-yl)-3,5-dihydrodibenzo[b,e]silin-10-yl)benzoate (BG-Sulfo646)* 7

*2.11. 4-((2-(2-((6-Chlorohexyl)oxy)ethoxy)ethyl)carbamoyl)-2-(5,5-dimethyl-3-(3-((2-sulfoethyl)carbamoyl)azetidin-1-ium-1-ylidene)-7-(3-((2-sulfoethyl)carbamoyl)azetidin-1-yl)-3,5-dihydrodibenzo[b,e]silin-10-yl)benzoate (Halo-Sulfo646)* 7

**3. Protein mass spectrometry** 8

**4. Cloning** 9

**5. Supplemental Figures** 10

**6. Supplemental Schemes** 14

**7. References** 15

### 1. General

All chemical reagents and anhydrous solvents for synthesis were purchased from commercial suppliers (Roth, Sigma-Aldrich, Fluka, Acros, Fluorochem, TCI) and were used without further purification or distillation. If necessary, solvents were degassed either by freeze-pump-thaw or by bubbling $N_2$ through the vigorously stirred solution for several minutes.

**BG-NH$_2$** and **CA-NH$_2$** were prepared and obtained as described previously.[1]

LC-MS was performed on a Shimadzu MS2020 connected to a Nexera UHPLC system equipped with a Waters ACQUITY UPLC BEH C18 (1.7 µm, 50 × 2.1 mm). Buffer A: 0.1% FA in $H_2O$ Buffer B: acetonitrile. The typical gradient was from 10% B for 0.5 min → gradient to 90% B over 4.5 min → 90% B for 0.5 min → gradient to 99% B over 0.5 min with 1 mL/min flow.

High resolution mass spectrometry was performed using a Bruker maXis II ETD hyphenated with a Shimadzu Nexera system. The instruments were controlled via Brukers otofControl 4.1 and Hystar 4.1 SR2 (4.1.31.1) software. The acquisition rate was set to 3 Hz and the following source parameters were used for positive mode electrospray ionization: End plate offset = 500 V; capillary voltage = 3800 V; nebulizer gas pressure = 45 psi; dry gas flow = 10 L/min; dry temperature = 250 °C. Transfer, quadrupole and collision cell settings are mass range dependent and were fine-adjusted with consideration of the respective analyte's molecular weight. For internal calibration sodium format clusters were used. Samples were desalted via fast liquid chromatography. A Supelco Titan™ C18 UHPLC Column, 1.9 µm, 80 Å pore size, 20 × 2.1 mm and a 2 min gradient from 10 to 98% aqueous MeCN with 0.1% FA ($H_2O$: Carl Roth GmbH + Co. KG ROTISOLV® Ultra LC-MS; MeCN: Merck KGaA LiChrosolv® Acetonitrile hypergrade for LC-MS; FA - Merck KGaA LiChropur® Formic acid 98%-100% for LC-MS) was used for separation. Sample dilution in 10% aqueous ACN (hyper grade) and injection volumes were chosen dependent of the analyte's ionization efficiency. Hence, on-column loadings resulted between 0.25–5.0 ng. Automated internal re-calibration and data analysis of the recorded spectra were performed with Bruker's DataAnalysis 4.4 SR1 software.

Semi-preparative RP-HPLC was performed on a Waters e2695 system equipped with a 2998 PDA detector for product collection (at 550 or 650 nm) on a Supelco Ascentis® C18 HPLC Column (5 µm, 250 × 4.6 mm). Buffer A: 0.1% TFA in $H_2O$ Buffer B: MeCN. The typical gradient was from 10% B for 5 min → gradient to 90% B over 30 min → 90% B for 5 min → gradient to 99% B over 5 min with 4 mL/min flow. All compounds were obtained in >95% purity according to LCMS.

Abbreviations: Boc: *tert*-butoxycarbonyl; DIPEA: *N,N*-diisopropylethylamine; DBU: 1,8-diazabicyclo[5.4.0]undec-7-ene; DCM: dichloromethane; DMSO: dimethylsulfoxide; FA: formic acid; Fmoc: fluorenylmethyloxycarbonyl; HBTU: (2-(1*H*-benzotriazol-1-yl)-1,1,3,3-tetramethyluronium hexafluorophosphate; TFA: trifluoroacetic acid; TSTU: *N,N,N′,N′*-Tetramethyl-*O*-(*N*-succinimidyl)uronium tetrafluoroborate.

### 2. Synthesis

#### 2.1. General procedure A for Buchwald-Hartwig coupling and deprotection

A flame-dried Schlenk flask was charged with bistriflate **1** (1.0 equiv.), $Pd_2(dba)_3$ (0.2 equiv.), XPhos (0.3 equiv., $Cs_2CO_3$ (4.8 equiv.), and dissolved in dry 1,4-dioxanes (345 equiv.) under an argon atmosphere. Methyl azetidine 3-carboxylate hydrochloride (3.0 equiv.) was added and the reaction mixture was heated to reflux over night. After cooling, MeOH and a spoon of Celite were added, all volatiles were removed *in vacuo*, and the resulting residue subjected to dry-loaded FCC purification (MeOH/DCM = 0/100 → 20/80). The fractions containing the rhodamines were pooled and taken up in aqueous 1 M LiOH:MeOH:THF (1:1:2) and allowed to incubate for 3 h before all volatiles were removed *in vacuo*, and the resulting residue subjected to RP-HPLC purification.

#### 2.2. General procedure B for taurine coupling and deprotection

A 4 mL dram vial was charged with acid **2** (1.0 equiv.) dissolved in DMF (2 mg/1 mL) and DIPEA (8.0 equiv.) and TSTU (2.8 equiv.) were combined before taurine (3.0 equiv.) was added in one portion. The reaction mixture was allowed to incubate for 1 h before it was quenched by addition of HOAc (10.0 equiv.) and 10 vol% water and subjected to RP-HPLC purification. The product containing fractions were pooled and lyophilized, and upon dryness redissolved in TFA:DCM (1/19) and incubated for 3 h at r.t. before all volatiles were removed *in vacuo* and the resulting solid was co-evaporated twice from toluene to obtain the taurine-coupled rhodamine with free 6-acid **3** sufficiently pure for the next step.

#### 2.3. General procedure C for peptide coupling with TSTU

A 4 mL dram vial was charged with acid **3** (1.0 equiv.) dissolved in DMF (2 mg/1 mL) and DIPEA (8.0 equiv.) and TSTU (1.4 equiv.) were combined before either **BG-NH$_2$** or **CA-NH$_2$** (1.5 equiv.) was added in one portion. The reaction mixture was allowed to incubate for 1 h before it was quenched by addition of HOAc (10.0 equiv.) and 10 vol% water and subjected to RP-HPLC purification. The product containing fractions were pooled and lyophilized to obtain the desired product as a colorful powder.

### 2.4. 4-(*tert*-Butoxycarbonyl)-2-(3-(3-carboxyazetidin-1-ium-1-ylidene)-6-(3-carboxyazetidin-1-yl)-3*H*-xanthen-9-yl)benzoate (2a)

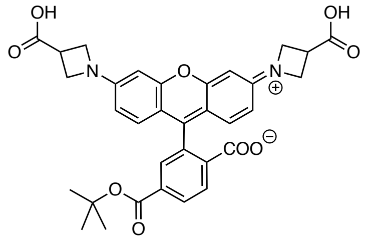

**2a** was prepared according to general procedure A.
bistriflate: **1a**

### 2.5. 4-Carboxy-2-(3-(3-((2-sulfoethyl)carbamoyl)azetidin-1-ium-1-ylidene)-6-(3-((2-sulfoethyl)carbamoyl)azetidin-1-yl)-3*H*-xanthen-9-yl)benzoate (3a)

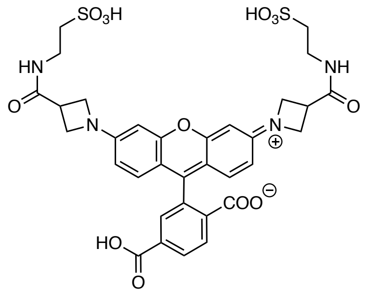

**3a** was prepared according to general procedure B.
acid: **2a**
**HRMS** (ESI): calc. for $C_{33}H_{33}N_4O_{13}S_2$ $[M+H]^+$: 757.1480, found: 757.1476.

### 2.6. 4-((4-(((2-Amino-9*H*-purin-6-yl)oxy)methyl)benzyl)carbamoyl)-2-(3-(3-((2-sulfoethyl)carbamoyl)azetidin-1-ium-1-ylidene)-6-(3-((2-sulfoethyl)carbamoyl)azetidin-1-yl)-3*H*-xanthen-9-yl)benzoate (BG-Sulfo549)

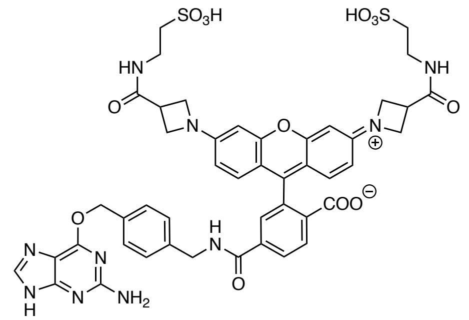

**BG-Sulfo549** was prepared according to general procedure C.
amine: **BG-NH₂**
acid: **3a**
**HRMS** (ESI): calc. for $C_{46}H_{46}N_{10}O_{13}S_2$ $[M+2H]^{2+}$: 505.1338, found: 505.1336.

### 2.7. 4-((2-(2-((6-Chlorohexyl)oxy)ethoxy)ethyl)carbamoyl)-2-(3-(3-((2-sulfoethyl)carbamoyl)azetidin-1-ium-1-ylidene)-6-(3-((2-sulfoethyl)carbamoyl)azetidin-1-yl)-3*H*-xanthen-9-yl)benzoate (Halo-Sulfo549)

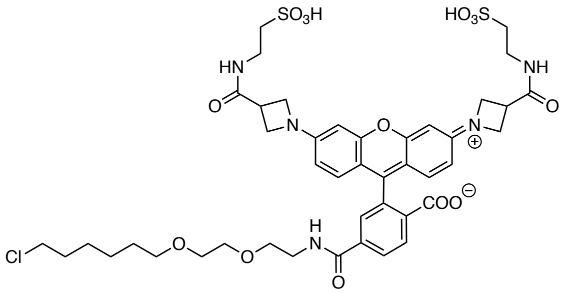


**Halo-Sulfo549** was prepared according to general procedure D.
amine: **Halo-NH$_2$**
acid: **3a**
**HRMS** (ESI): calc. for $C_{43}H_{53}ClN_5O_{14}S_2$ $[M+H]^+$: 962.2713, found: 962.2721.

### 2.8. 4-(*tert*-Butoxycarbonyl)-2-(3-(3-carboxyazetidin-1-ium-1-ylidene)-7-(3-carboxyazetidin-1-yl)-5,5-dimethyl-3,5-dihydrodibenzo[*b*,*e*]silin-10-yl)benzoate (2c)

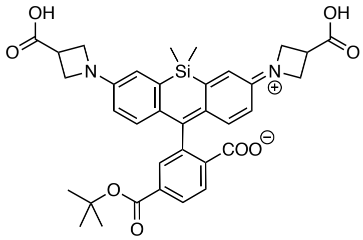


**2c** was prepared according to general procedure A.
bistriflate: **1c**
**HRMS** (ESI): calc. for $C_{35}H_{37}N_2O_8Si$ $[M+H]^+$: 641.2314, found: 641.2314.

### 2.9. 4-Carboxy-2-(5,5-dimethyl-3-(3-((2-sulfoethyl)carbamoyl)azetidin-1-ium-1-ylidene)-7-(3-((2-sulfoethyl)carbamoyl)azetidin-1-yl)-3,5-dihydrodibenzo[*b*,*e*]silin-10-yl)benzoate (3c)

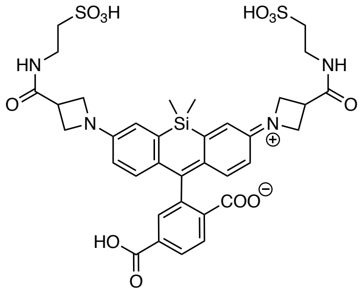


**3c** was prepared according to general procedure B.
acid: **2c**

#### 2.10. 4-((4-(2-(2-Amino-9*H*-purin-6-yl)ethyl)benzyl)carbamoyl)-2-(5,5-dimethyl-3-(3-((2-sulfoethyl)carbamoyl)azetidin-1-ium-1-ylidene)-7-(3-((2-sulfoethyl)carbamoyl)azetidin-1-yl)-3,5-dihydrodibenzo[*b*,*e*]silin-10-yl)benzoate (BG-Sulfo646)

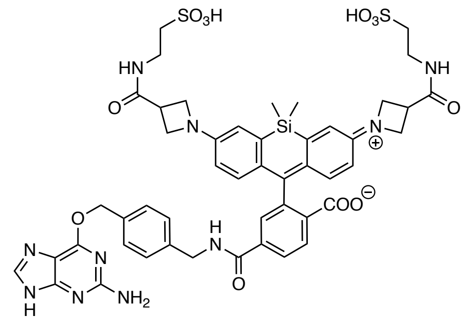


**BG-Sulfo646** was prepared according to general procedure C.
amine: **BG-NH₂**
acid: **3c**
**HRMS** (ESI): calc. for $C_{48}H_{52}N_{10}O_{12}S_2Si$ $[M+2H]^{2+}$: 526.1483, found: 526.1481.

#### 2.11. 4-((2-(2-((6-Chlorohexyl)oxy)ethoxy)ethyl)carbamoyl)-2-(5,5-dimethyl-3-(3-((2-sulfoethyl)carbamoyl)azetidin-1-ium-1-ylidene)-7-(3-((2-sulfoethyl)carbamoyl)azetidin-1-yl)-3,5-dihydrodibenzo[*b*,*e*]silin-10-yl)benzoate (Halo-Sulfo646)

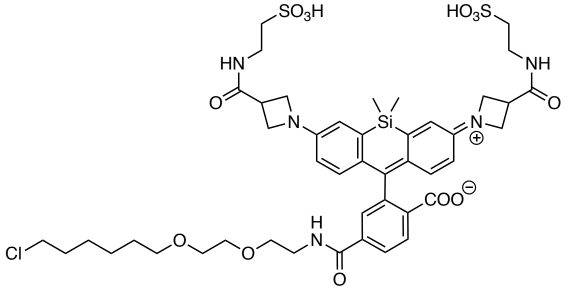


**Halo-Sulfo646** was prepared according to general procedure D.
amine: **Halo-NH₂**
acid: **3c**
**HRMS** (ESI): calc. for $C_{45}H_{56}ClN_5O_{13}S_2Si$ $[M-2H]^{2-}$: 500.6392, found: 500.6390.

### 3. Protein mass spectrometry

SNAP$_f$ sequence:
MASWSHPQFEKGADDDDKVPHMDKDCEMKRTTLDSPLGKLELSGCEQGLHRIIFLGKGTSAADAV
EVPAPAAVLGGPEPLMQATAWLNAYFHQPEAIEEFPVPALHHPVFQQESFTRQVLWKLLKVVKFG
EVISYSHLAALAGNPAATAAVKTALSGNPVPILIPCHRVVQGDLDVGGYEGGLAVKEWLLAHEGH
RLGKPGLGAPGFSSISAHHHHHHHHHH

Strep-Tag II, Enterokinase-site, SNAPf, His-Tag

After His-tag purification, two posttranslational SNAP$_f$ constructs were observed: one with removed start codon SNAP$_f$ (2–222) and one without the N-terminal Strep-Tag II and Enterokinase-site SNAP$_f$ (22–222):

| | calc. | found | Δ ppm |
|---|---|---|---|
| SNAP$_f$ (2–222) | 23865.101 | 23865.141 | 1.67 |
| SNAP$_f$ (22–222) | 21618.103 | 21618.142 | 1.80 |
| SNAP$_f$ (2–222):JF$_{549}$ | 24420.317 | 24420.357 | 1.65 |
| SNAP$_f$ (22–222): JF$_{549}$ | 22173.319 | 22173.349 | 1.36 |
| SNAP$_f$ (2–222): JF$_{646}$ | 24462.386 | 24462.386 | 0.00 |
| SNAP$_f$ (22–222): JF$_{646}$ | 22215.358 | 22215.386 | 1.72 |
| SNAP$_f$ (2–222):Sulfo549 | 24722.305 | 24722.340 | 1.43 |
| SNAP$_f$ (22–222):Sulfo549 | 22475.307 | 22475.337 | 1.35 |
| SNAP$_f$ (2–222):Sulfo646 | 24764.334 | 24764.387 | 2.16 |
| SNAP$_f$ (22–222):Sulfo646 | 22517.336 | 22517.387 | 2.28 |

Halo sequence:
MHHHHHHHHHHENLYFQGIGTGFPFDPHYVEVLGERMHYVDVGPRDGTPVLFLHGNPTSSYVWRN
IIPHVAPTHRCIAPDLIGMGKSDKPDLGYFFDDHVRFMDAFIEALGLEEVVLVIHDWGSALGFHW
AKRNPERVKGIAFMEFIRPIPTWDEWPEFARETFQAFRTTDVGRKLIIDQNVFIEGTLPMGVVRP
LTEVEMDHYREPFLNPVDREPLWRFPNELPIAGEPANIVALVEEYMDWLHQSPVPKLLFWGTPGV
LIPPAEAARLAKSLPNCKAVDIGPGLNLLQEDNPDLIGSEIARWLSTLEI

Halo, His-Tag

| | calc. | found | Δ ppm |
|---|---|---|---|
| Halo | 35465.901 | 35465.985 | 2.37 |
| Halo:JF$_{549}$ | 36089.251 | 36089.293 | 1.16 |
| Halo:JF$_{646}$ | 36131.230 | 36131.297 | 1.85 |
| Halo:Sulfo549 | 36391.188 | 36391.291 | 2.83 |
| Halo: Sulfo646 | 36433.217 | 36433.317 | 2.74 |

### 4. Cloning

SNAP–TM–Halo sequence:

METDTLLLWVLLLWVPGSTGDYPYDVPDYAGAQPARSMDKDCEMKRTTLDSPLGKLELSGCEQGL
HEIIFLGKGTSAADAVEVPAPAAVLGGPEPLMQATAWLNAYFHQPEAIEEFPVPALHHPVFQQES
FTRQVLWKLLKVVKFGEVISYSHLAALAGNPAATAAVKTALSGNPVPILIPCHRVVQGDLDVGGY
EGGLAVKEWLLAHEGHRLGKPGLGGRLEVLFQGVDEQKLISEEDLNAVGQDTQEVIVVPHSLPFK
VVVISAILALVVLTIISLIILIMLWQKKPRGAQPARSGSEIGTGFPFDPHYVEVLGERMHYVDVG
PRDGTPVLFLHGNPTSSYVWRNIIPHVAPTHRCIAPDLIGMGKSDKPDLGYFFDDHVRFMDAFIE
ALGLEEVVLVIHDWGSALGFHWAKRNPERVKGIAFMEFIRPIPTWDEWPEFARETFQAFRTTDVG
RKLIIDQNVFIEGTLPMGVVRPLTEVEMDHYREPFLNPVDREPLWRFPNELPIAGEPANIVALVE
EYMDWLHQSPVPKLLFWGTPGVLIPPAEAARLAKSLPNCKAVDIGPGLNLLQEDNPDLIGSEIAR
WLSTLEISGVD*

IgK signal peptide, HA-tag, myc, SNAP, transmembrane domain, Halo-Tag

Halo–TM–SNAP sequence:

METDTLLLWVLLLWVPGSTGDYPYDVPDYAGAQPARSGSEIGTGFPFDPHYVEVLGERMHYVDVG
PRDGTPVLFLHGNPTSSYVWRNIIPHVAPTHRCIAPDLIGMGKSDKPDLGYFFDDHVRFMDAFIE
ALGLEEVVLVIHDWGSALGFHWAKRNPERVKGIAFMEFIRPIPTWDEWPEFARETFQAFRTTDVG
RKLIIDQNVFIEGTLPMGVVRPLTEVEMDHYREPFLNPVDREPLWRFPNELPIAGEPANIVALVE
EYMDWLHQSPVPKLLFWGTPGVLIPPAEAARLAKSLPNCKAVDIGPGLNLLQEDNPDLIGSEIAR
WLSTLEISGVDEQKLISEEDLNAVGQDTQEVIVVPHSLPFKVVVISAILALVVLTIISLIILIML
WQKKPRGAQPARSMDKDCEMKRTTLDSPLGKLELSGCEQGLHEIIFLGKGTSAADAVEVPAPAAV
LGGPEPLMQATAWLNAYFHQPEAIEEFPVPALHHPVFQQESFTRQVLWKLLKVVKFGEVISYSHL
AALAGNPAATAAVKTALSGNPVPILIPCHRVVQGDLDVGGYEGGLAVKEWLLAHEGHRLGKPGLG
GRLEVLFQGVD*

IgK signal peptide, HA-tag, Halo-Tag, myc, transmembrane domain, SNAP

Halo-GLP1R sequence:

MALPVTALLLPLALLLHAARPAAASGIDYKDDDDKAGIDAIGSEIGTGFPFDPHYVEVLGERMHY
VDVGPRDGTPVLFLHGNPTSSYVWRNIIPHVAPTHRCIAPDLIGMGKSDKPDLGYFFDDHVRFMD
AFIEALGLEEVVLVIHDWGSALGFHWAKRNPERVKGIAFMEFIRPIPTWDEWPEFARETFQAFRT
TDVGRKLIIDQNVFIEGTLPMGVVRPLTEVEMDHYREPFLNPVDREPLWRFPNELPIAGEPANIV
ALVEEYMDWLHQSPVPKLLFWGTPGVLIPPAEAARLAKSLPNCKAVDIGPGLNLLQEDNPDLIGS
EIARWLSTLEISGDIRPQGATVSLWETVQKWREYRRQCQRSLTEDPPPATDLFCNRTFDEYACWP
DGEPGSFVNVSCPWYLPWASSVPQGHVYRFCTAEGLWLQKDNSSLPWRDLSECEESKRGERSSPE
EQLLFLYIIYTVGYALSFSALVIASAILLGFRHLHCTRNYIHLNLFASFILRALSVFIKDAALKW
MYSTAAQQHQWDGLLSYQDSLSCRLVFLLMQYCVAANYYWLLVEGVYLYTLLAFSVLSEQWIFRL
YVSIGWGVPLLFVVPWGIVKYLYEDEGCWTRNSNMNYWLIIRLPILFAIGVNFLIFVRVICIVVS
KLKANLMCKTDIKCRLAKSTLTLIPLLGTHEVIFAFVMDEHARGTLRFIKLFTELSFTSFQGLMV
AILYCFVNNEVQLEFRKSWERWRLEHLHIQRDSSMKPLKCPTSSLSSGATAGSSMYTATCQASCS
*

CD8 signal peptide, FLAG-tag, Halo-tag, GLP1R

### 5. Supplemental Figures

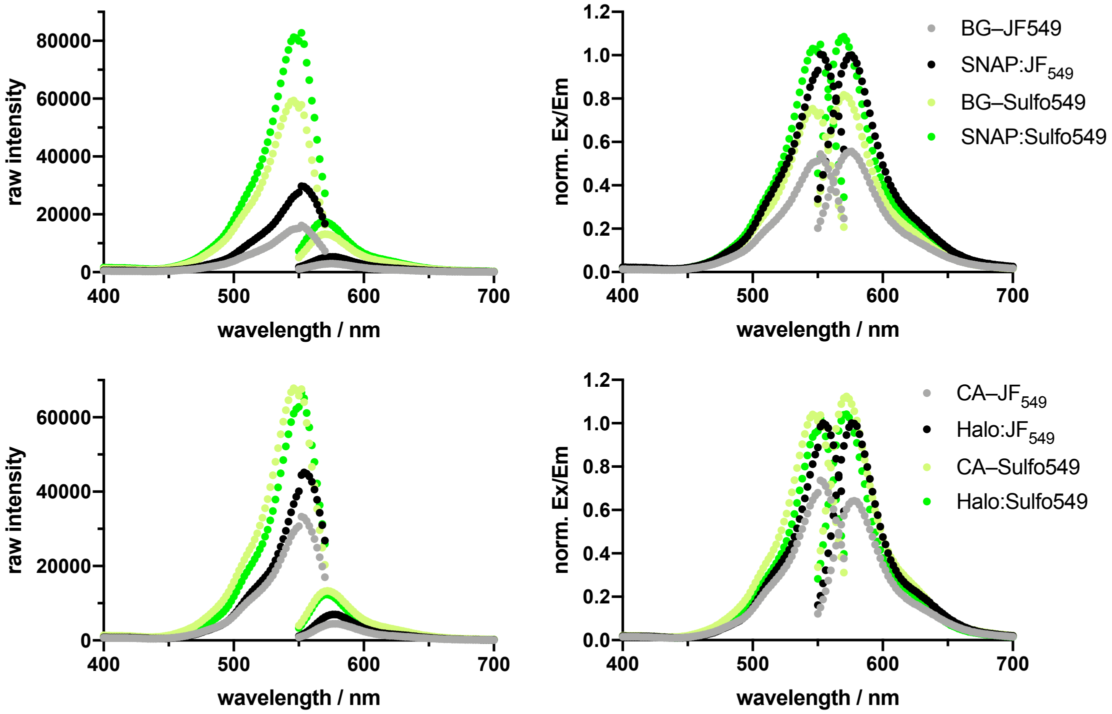


**Supplemental Figure 1: Excitation and emission spectra of bound and unbound 549 probes with raw intensity (left) and normalized to JF dye excitation and emission (right, cf. Figure 2).**

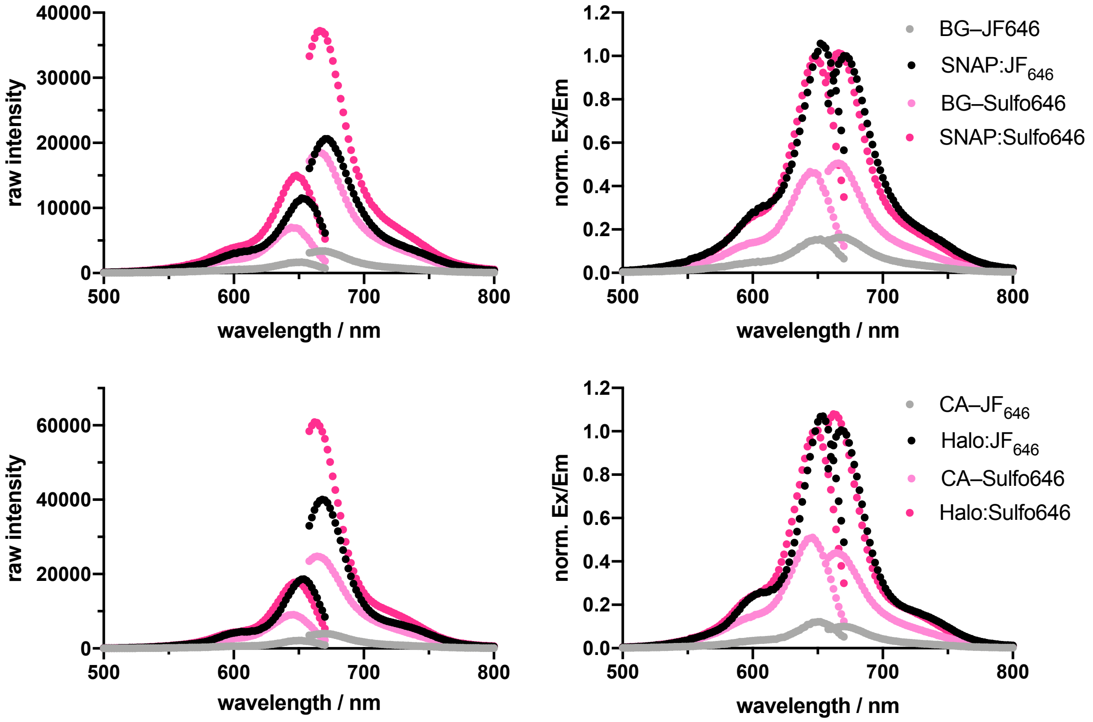


**Supplemental Figure 2: Excitation and emission spectra of bound and unbound 646 probes with raw intensity (left) and normalized to JF dye excitation and emission (right, cf. Figure 2).**

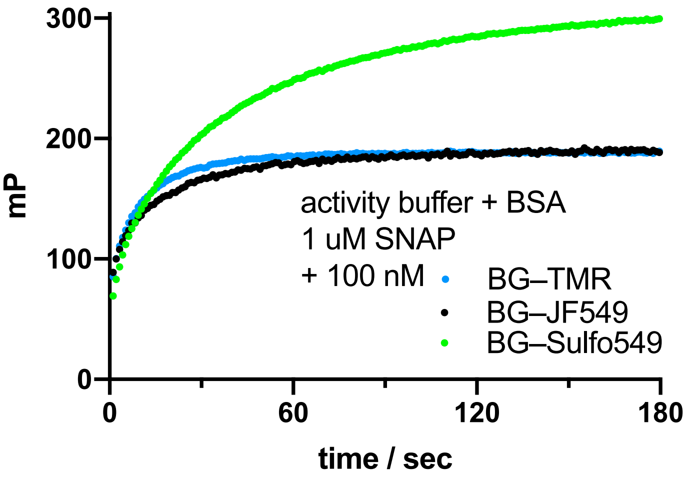


**Supplemental Figure 3: Reaction kinetics of BG–TMR, BG–JF$_{549}$ and BG–Sulfo549 towards SNAP-tag measured by fluorescence polarization.**

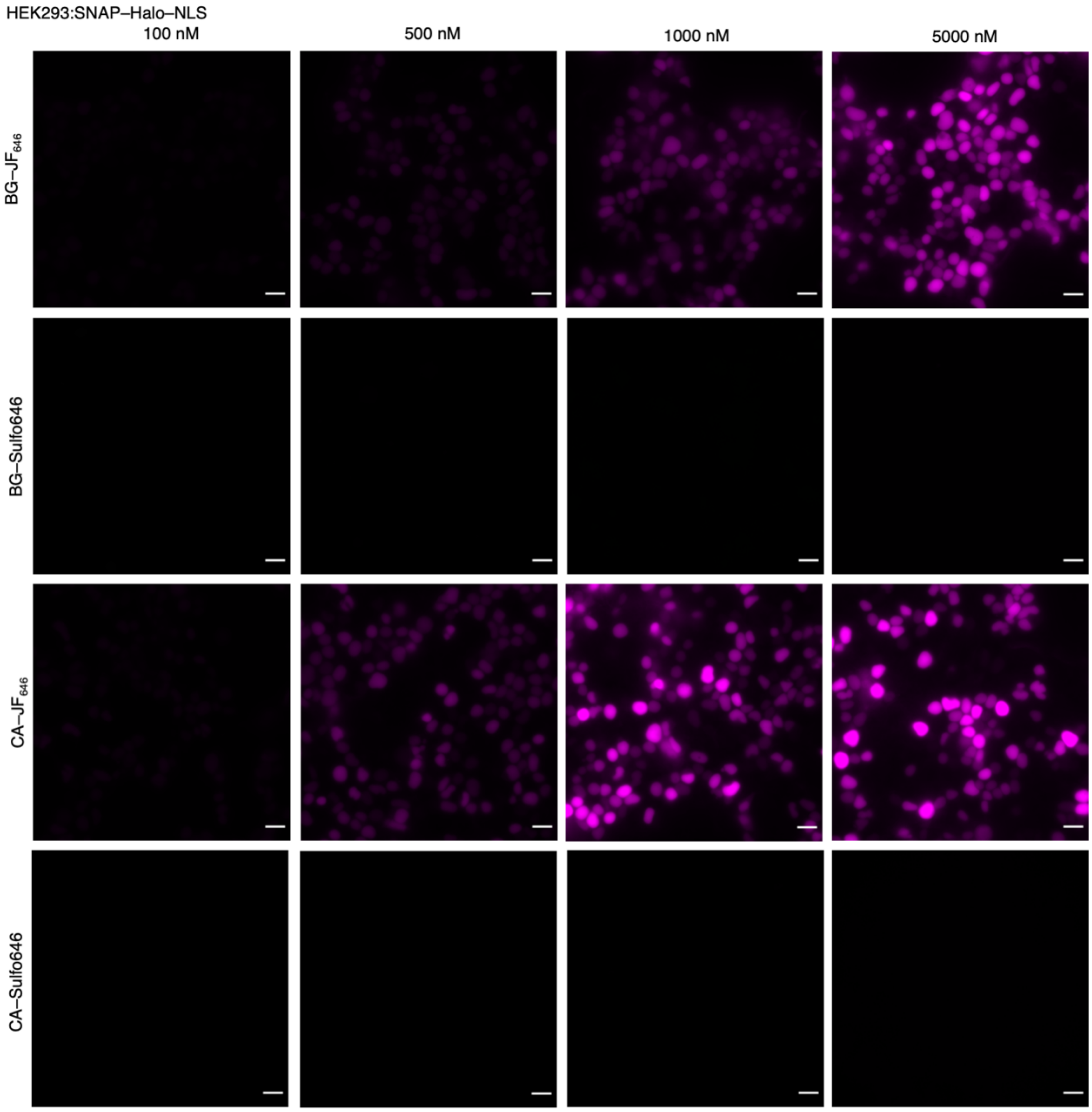


**Supplemental Figure 4: Titration of dyes in SNAP–Halo–NLS transfected HEK293 cells.** SNAP–Halo–NLS transfected HEK293 cells were treated with 100, 500, 1000 and 5000 nM of BG–$JF_{646}$, BG–Sulfo646, CA–$JF_{646}$ or CA–Sulfo646.

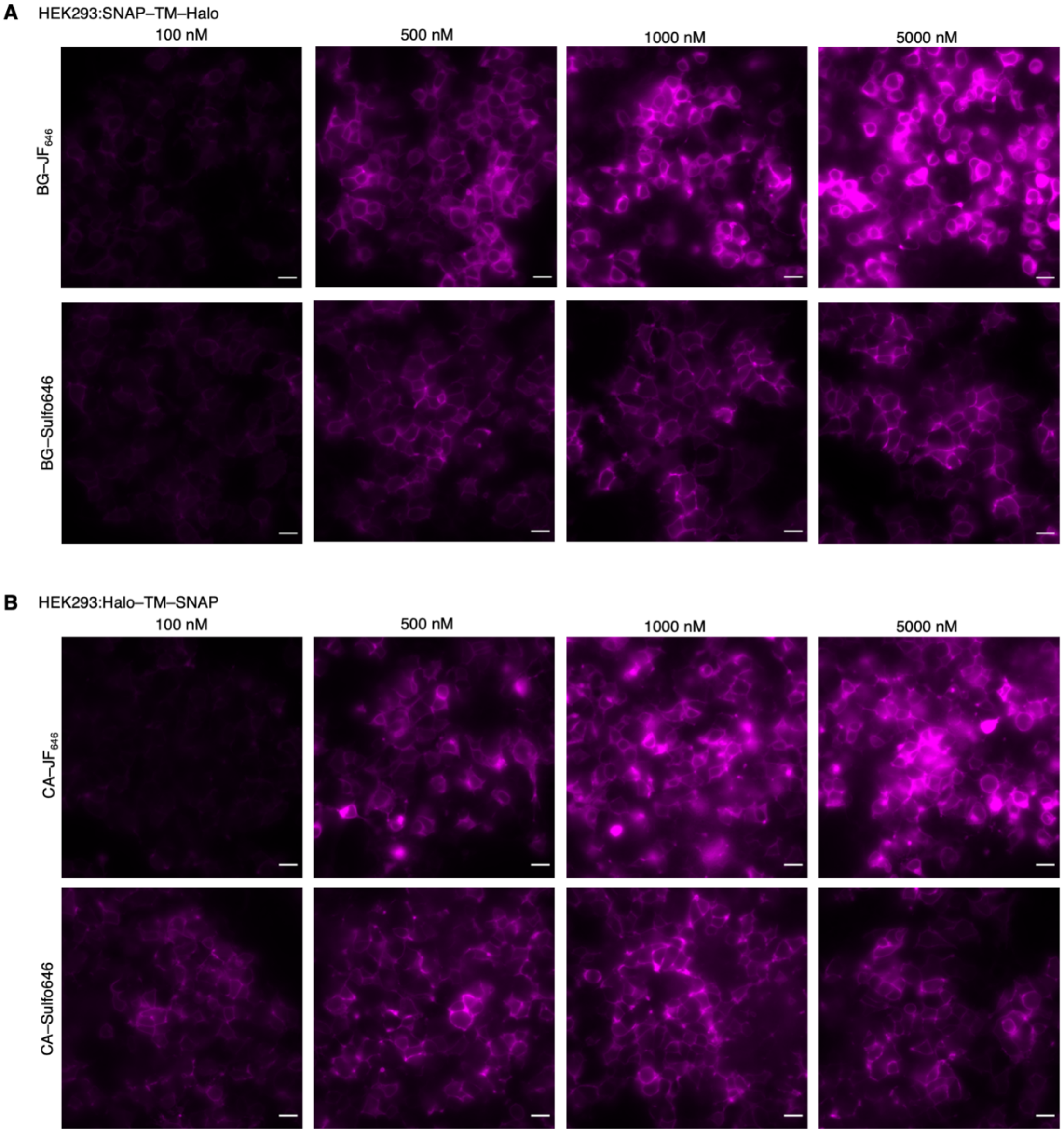


**Supplemental Figure 5: Titration of dyes in SNAP–Halo transfected HEK293 cells. A)** SNAP–TM–Halo transfected HEK293 cells were treated with 100, 500, 1000 and 5000 nM of BG–JF646 or BG–Sulfo646. **A)** Halo–TM–SNAP transfected Hek293 cells were treated with 100, 500, 1000 and 5000 nM of CA–JF646 or CA–Sulfo646.

### 6. Supplemental Schemes

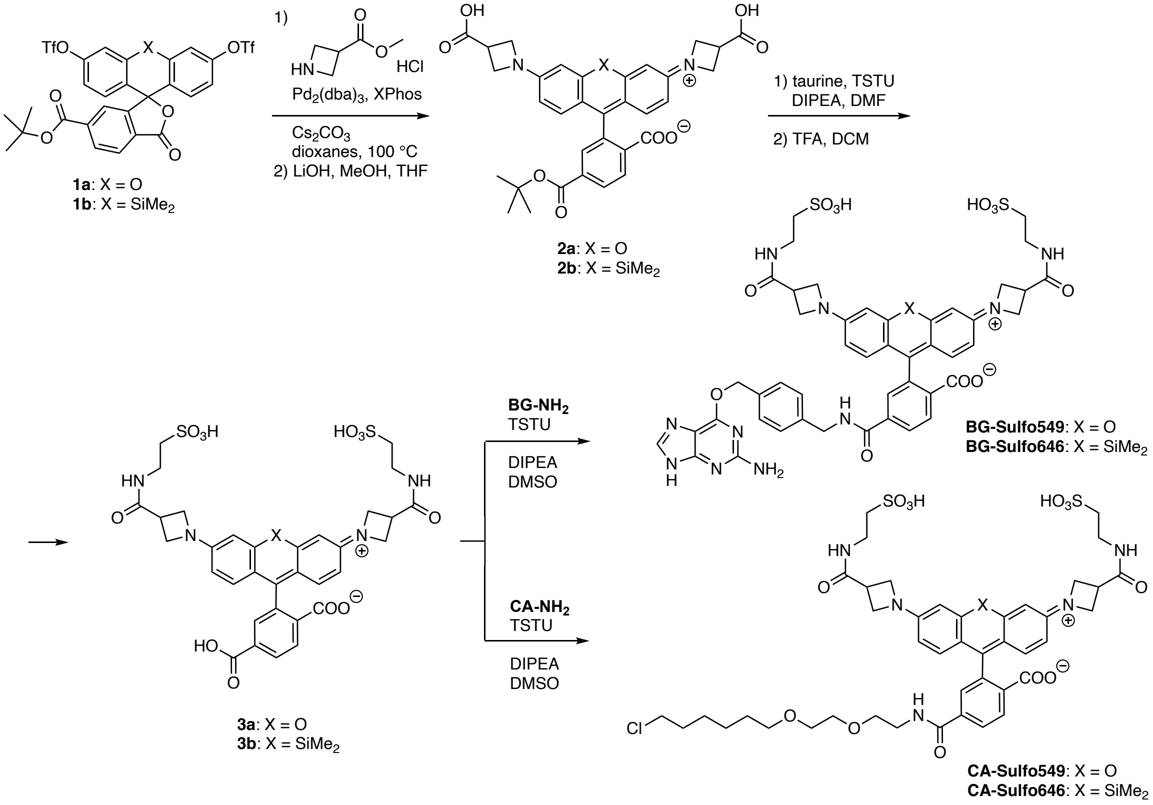


**Supplemental Scheme 1: Synthesis of Sulfo dyes.**
